# The cell-type underpinnings of regional brain entropy (BEN) in human brain

**DOI:** 10.1101/2024.11.22.624840

**Authors:** Donghui Song

## Abstract

Brain entropy (BEN) indicates the irregularity, unpredictability and complexity of brain activity. Research using regional brain entropy (rBEN) based on fMRI has established its associations with structural and functional brain networks, as well as their coupling. These relationships are influenced by the sensorimotor-association (S-A) axis and cytoarchitectural organization. Recent studies have also revealed links between functional connectome gradients and cell-type, suggesting that rBEN may have a biological basis at the cellular level. However, this possibility remains unexplored at specifically cellular level.

In this study, we analyzed mean rBEN maps derived from 176 participants using HCP 7T data and correlated these with publicly available cell-type datasets. Our findings rBEN exhibited a positive correlation with oligodendrocytes (Oligo) and a negative correlation with parvalbumin-positive interneurons (PVALB) under both resting-state and movie-watching conditions at the whole-brain level. Furthermore, we observed that the S-A axis modulates the relationship between rBEN and cell-type. Specifically, L5 extratelencephalic neurons (L5 ET) showed a positive correlation with rBEN in unimodal cortex but a negative correlation in multimodal cortex and interneurons somatostatin (SST) was correlated with rBEN only in multimodal cortex.

These results provide further evidence that rBEN has a biological foundation at the cellular level, with spatial heterogeneity in its associations with different cell types. Moreover, rBEN appears to capture information beyond functional connectivity network.

## 1. Introduction

The brain entropy (BEN) indicates the irregularity, disorder, uncertainty, and complexity of human brain activity. Studies using functional magnetic resonance imaging (fMRI)-based regional BEN (rBEN) have identified rBEN contrasts between gray and white matter (Wang, Li et al. 2014), establishing its structural foundation relevance (Del Mauro and Wang 2024, Song and Wang 2024) as well as its relationship with neurochemical signals, including neurotransmitters(Song and Wang 2024) and hormones (Song and Wang 2024). Furthermore, the relationship between rBEN and both structural and functional networks has been characterized (Song 2024). At the same time, recent research has revealed cell-type underpinnings of the human functional cortical connectome that differentially enriched cells follow the spatial topography of both functional gradients and associated large-scale networks (Zhang, Anderson et al. 2023). Our recent research has revealed that functional connectomes and connectome gradients are associated with rBEN and that these relationships are influenced by cytoarchitectural organization. The sensorimotor-association (S-A) axis also plays a vital role in the relationships between rBEN, brain networks, and cytoarchitectural organization. These findings suggest a potential link between rBEN distribution and specific cell types. However, the cellular basis of BEN remains unexplored. This study aims to bridge the gap between cellular types and macroscopic brain activity, offering novel insights into the biological foundations of rBEN.

## 2. Methods

### 2.1 Dataset

The rBEN maps were calculated from preprocessed resting-state fMRI and movie-watching fMRI from HCP 7T release (Van Essen, Smith et al. 2013). These rBEN maps used in this study are the same as that in our previous research (Song and Wang 2024, Song 2024, Song and Wang 2024, Song and Wang 2024), with a total of 176 participants included. For each participant, the resting-state rBEN (rsBEN) and movie-watching rBEN (mvBEN) were segmented into 400 parcels based on the Schaefer 400 (Schaefer, Kong et al. 2018). For details on the calculation and parameter selection of rBEN, please refer to (Wang, Li et al. 2014, Wang 2021, Song and Wang 2024).

The archetypal sensorimotor-associationaxis from https://github.com/PennLINC/S-A_ArchetypalAxis (Sydnor, Larsen et al. 2021).

The processed cell-type data were obtained from https://github.com/XihanZhang/human-cellular-func-con (Zhang, Anderson et al. 2023). The raw cell-type data originated from single-nucleus droplet-based sequencing (snDrop-seq) data available on the cellxgene-census platform (https://cellxgene.cziscience.com/collections/d17249d2-0e6e-4500-abb8-e6c93fa1ac6f) (Jorstad, Close et al. 2023) and the Gene Expression Omnibus website (GSE97930; https://www.ncbi.nlm.nih.gov/geo) (Lake, Chen et al. 2018). In brief, 24 cell types from Jorstad dataset, 18 cell types were derived from visual (Lake dataset VIS) and frontal (Lake dataset DFC) samples separately from Lake dataset. The cell-type abundances for each AHBA cortical sample were mapped to the cortical vertices represented in fsaverage6 surface space and then parceled into the Schaefer 400 atlas (Zhang, Anderson et al. 2023).

## 3. Results

We removed the parcels without cell data, leaving a total of 339 parcels with data. Then, spatial correlation was calculated between rsBEN, mvBEN in whole brain and in the unimodal (S-A < 201) and multimodal (S-A > 200) cortices along the S-A axis, using the Pearson correlation coefficient for each with the 24 cellular classes with distinct laminar specialization, developmental origins, morphology, spiking patterns and broad projection targets from (Jorstad, Close et al. 2023). The nine GABAergic inhibitory interneurons (PAX6, SNCG, VIP, LAMP5, LAMP5 LHX6, Chandelier, PVALB, SST CHODL and SST), nine glutamatergic excitatory neurons (L2/3 IT, L4 IT, L5 IT, L6 IT, L5 ET, L5/6 NP, L6 CT, L6b and L6 IT Car3), and six non-neuronal cells (Astro, Endo, VLMC, Oligo, OPC and Micro/PVM).

We define a correlation coefficient greater than 0.3 as indicating a significant correlation. The corresponding p-values are provided for the whole brain correlation (n=339, *p*<1.44 ×10^−8^), unimodal region (n=162, *p*<0.001), multimodal cortex (n=177, *p*<0.001), and the difference between multimodal and unimodal cortices (*p*<0.005).

The results (Fig.1) show that rsBEN is positively correlated with oligodendrocytes (Oligo) (rsBEN: R_n=339_=0.30, mvBEN: R_n=339_=0.36) and negatively correlated with parvalbumin (PVALB) (rsBEN: R_n=339_=0.30, mvBEN: R_n=339_=0.38), both in the resting-state and movie-watching conditions. We also observed that the relationship between rsBEN and cell-type is influenced by the S-A axis. Specifically, these cells primarily include layer 5 extratelencephalic (L5 ET) and interneurons somatostatin (SST). The correlation between rsBEN and L5 ET shows opposite patterns across unimodal and multimodal cortex: it is positively correlated in unimodal cortex (rsBEN: R_n=162_=0.2, mvBEN: R_n=162_=0.31) and negatively correlated in multimodal cortex (rsBEN: R_n=177_=-0.2, mvBEN: R_n=177_=-0.17). In contrast, the positive correlation between rsBEN and SST is observed in multimodal cortex (rsBEN: R_n=177_=0.32, mvBEN: R_n=177_=0.31) but not in unimodal cortex. The results from (Lake, Chen et al. 2018) are presented in sFig1 for Lake dfc and sFig2 for Lake vis.

**Fig. 1.**
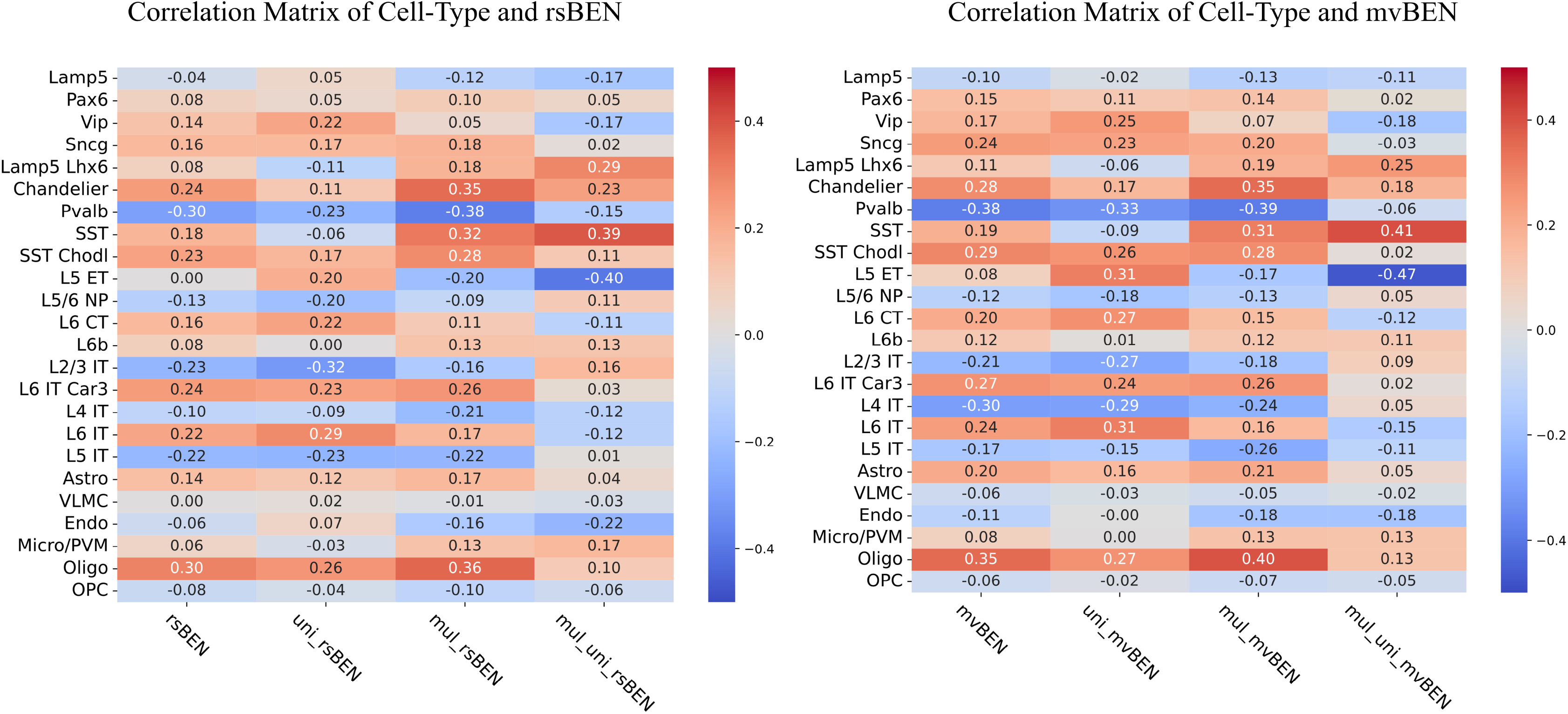
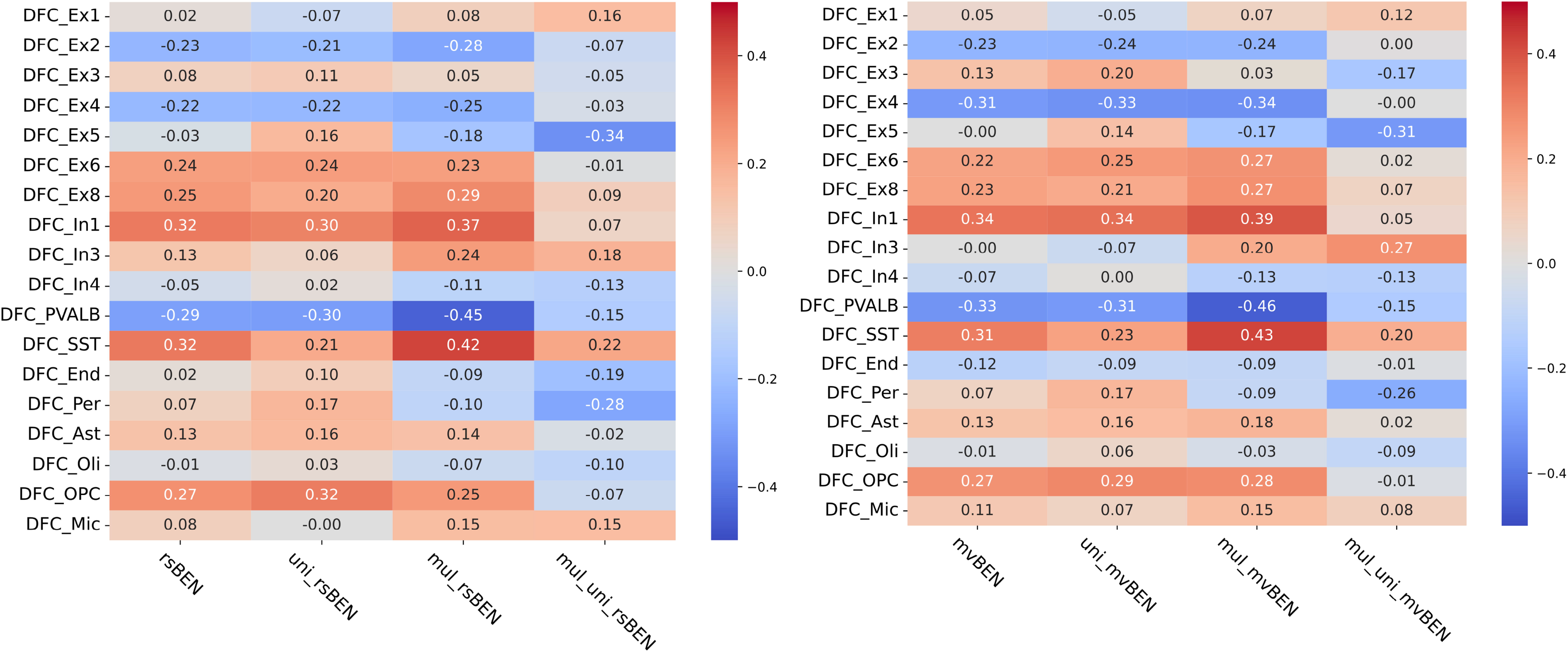
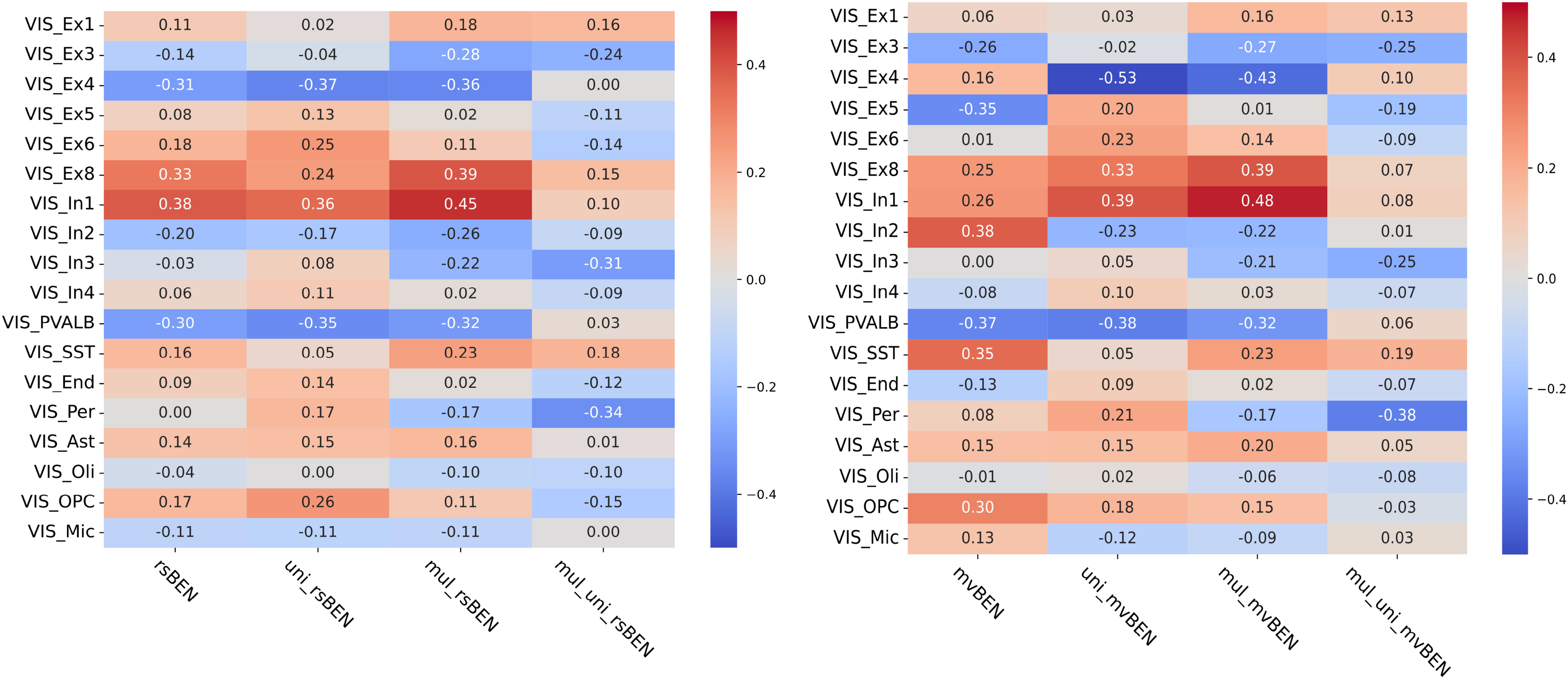
Correlation between cell-type and rBEN. The left panel shows the correlation between cell-type and rsBEN, while the right panel displays the correlation with mvBEN. The y-axis represents the cell-types, and the x-axis shows the correlations across the whole brain, unimodal cortex, multimodal cortex, and the difference in correlation between unimodal and multimodal cortices. The color bar represents the strength of the correlation coefficient, with warm colors indicating positive correlations and cool colors indicating negative correlations. The correlation coefficients are also displayed within the figures.

## 4. Discussion

This study is the first to demonstrate that rBEN has a biological basis rooted in cell types. The spatial distribution of rBEN is primarily positively correlated with the spatial enrichment of Oligo and negatively correlated with the distribution of PVALB. We also found that along S-A axis, the enrichment of cell types in unimodal cortex and multimodal cortex significantly influences rBEN. Specifically, L5 ET and SST play distinct roles in these regions: in unimodal regions, L5 ET shows a positive correlation with rBEN, whereas in multimodal regions, L5 ET exhibits a negative correlation with rBEN. SST, on the other hand, is positively correlated with rBEN in multimodal cortex but shows no significant correlation in unimodal cortex.

Oligo are glial cells responsible for producing the myelin sheath, which accelerates the transmission of electrical signals between neurons (Emery 2010, Simons and Nave 2016, Moore, Meschkat et al. 2020). The positive correlation between Oligo and rBEN may be related to enhanced information reception and transmission. Our previous research found a positive association between the local efficiency of high-frequency MEG functional networks and the coupling between BOLD-network and MEG-network (Song 2024). An increase in rBEN suggests that the efficiency of information generation and transmission outweighs the loss of efficiency, aligning with the role of Oligo in improving the efficiency of information transfer between neurons. PVALB, on the other hand, is an inhibitory interneuron that plays a crucial role in regulating network dynamics and generating oscillations in the brain (Sohal, Zhang et al. 2009, Kim, Ährlund-Richter et al. 2016). Its negative correlation with rBEN may indicate that PVALB promotes information processing by regulating regional network synchronization and rhythmicity, thereby reducing entropy. L5 ET neurons are a subclass of pyramidal neurons in the neocortex that transmit signals from the cerebral cortex to other parts of the brain and body (Kalmbach, Hodge et al. 2021, Moberg and Takahashi 2022). In unimodal cortex, L5 ET cells show a positive correlation with rBEN, indicating that they facilitate information generation and transmission. In multimodal cortex, L5 ET shows a negative correlation with rBEN, suggesting that they play a role in complex cognitive processing by helping to enhance information processing efficiency. SST is inhibitory GABAergic interneurons that suppress the activity of target neurons by releasing GABA (gamma-aminobutyric acid) that regulate inhibitory control within neural networks, thereby influencing the integration and propagation of information (Urban-Ciecko and Barth 2016, Song, Yoon et al. 2021, Mòdol, Moissidis et al. 2024). As SST cells are primarily enriched in multimodal cortices, previous studies have found a negative correlation between resting-state rBEN in the fronto-parietal network and fluid intelligence and education levels(Wang 2021, Del Mauro and Wang 2024). The SST was also found to be significantly positively correlated with the principal cortical functional gradient (Zhang, Anderson et al. 2023). However, our results differ in terms of correlation with other cell types compared to studies based on functional connectome gradients. Combined with our previous research on the relationship between rBEN and brain networks (Song 2024, Song and Wang 2024, Song and Wang 2024), these findings suggest that rBEN may capture information beyond network connectivity.

As a preliminary study and an initial draft, further research is needed, such as conducting more rigorous statistical tests, examining the overall contribution of cell-type to rBEN, and determining whether cell-type can distinguish the distribution patterns of rBEN. In summary, the study provides an initial conclusion that the spatial distribution of cell-type is closely related to rBEN, suggesting that rBEN has a cellular-level biological basis.

## Acknowledgments

MRI data were provided by the Human Connectome Project, WU-Minn Consortium (Principal Investigators: David Van Essen and Kamil Ugurbil; 1U54MH091657) funded by the 16 NIH Institutes and Centers that support the NIH Blueprint for Neuroscience Research; and by the McDonnell Center for Systems Neuroscience at Washington University. We thank Xihan Zhang et al (Zhang, Anderson et al. 2023) for releasing their dataset.

## References

Del Mauro, G. and Z. Wang (2024). “Associations of brain entropy estimated by resting state fMRI with physiological indices, body mass index, and cognition.” Journal of Magnetic Resonance Imaging 59(5): 1697–1707.

Del Mauro, G. and Z. Wang (2024). “rsfMRI-based Brain Entropy is negatively correlated with Gray Matter Volume and Surface Area.” bioRxiv: 2024.2004. 2028.591371.

Emery, B. (2010). “Regulation of oligodendrocyte differentiation and myelination.” Science 330(6005): 779–782.

Jorstad, N. L., J. Close, N. Johansen, A. M. Yanny, E. R. Barkan, K. J. Travaglini, D. Bertagnolli, J. Campos, T. Casper and K. Crichton (2023). “Transcriptomic cytoarchitecture reveals principles of human neocortex organization.” Science 382(6667): eadf6812.

Kalmbach, B. E., R. D. Hodge, N. L. Jorstad, S. Owen, R. de Frates, A. M. Yanny, R. Dalley, M. Mallory, L. T. Graybuck and C. Radaelli (2021). “Signature morpho-electric, transcriptomic, and dendritic properties of human layer 5 neocortical pyramidal neurons.” Neuron 109(18): 2914-2927. e2915.

Kim, H., S. Ährlund-Richter, X. Wang, K. Deisseroth and M. Carlén (2016). “Prefrontal parvalbumin neurons in control of attention.” Cell 164(1): 208–218.

Lake, B. B., S. Chen, B. C. Sos, J. Fan, G. E. Kaeser, Y. C. Yung, T. E. Duong, D. Gao, J. Chun and P. V. Kharchenko (2018). “Integrative single-cell analysis of transcriptional and epigenetic states in the human adult brain.” Nature biotechnology 36(1): 70–80.

Moberg, S. and N. Takahashi (2022). “Neocortical layer 5 subclasses: from cellular properties to roles in behavior.” Frontiers in Synaptic Neuroscience 14: 1006773.

Mòdol, L., M. Moissidis, M. Selten, F. Oozeer and O. Marín (2024). “Somatostatin interneurons control the timing of developmental desynchronization in cortical networks.” Neuron.

Moore, S., M. Meschkat, T. Ruhwedel, A. Trevisiol, I. D. Tzvetanova, A. Battefeld, K. Kusch, M. H. Kole, N. Strenzke and W. Möbius (2020). “A role of oligodendrocytes in information processing.” Nature communications 11(1): 5497.

Schaefer, A., R. Kong, E. M. Gordon, T. O. Laumann, X.-N. Zuo, A. J. Holmes, S. B. Eickhoff and B. T. Yeo (2018). “Local-global parcellation of the human cerebral cortex from intrinsic functional connectivity MRI.” Cerebral cortex 28(9): 3095–3114.

Simons, M. and K.-A. Nave (2016). “Oligodendrocytes: myelination and axonal support.” Cold Spring Harbor perspectives in biology 8(1): a020479.

Sohal, V. S., F. Zhang, O. Yizhar and K. Deisseroth (2009). “Parvalbumin neurons and gamma rhythms enhance cortical circuit performance.” Nature 459(7247): 698–702.

Song, D.-H. and Z. Wang (2024). “Regional Brain Entropy During Movie-watching.” bioRxiv: 2024.2006. 2012.598767.

Song, D.-H. and Z. Wang (2024). “The Relationships of Resting-state Brain Entropy (BEN), Ovarian Hormones and Behavioral Inhibition and Activation Systems (BIS/BAS).” bioRxiv: 2024.2006. 2004.595915.

Song, D. (2024). “Regional Brain Entropy, Brain Network and Structural-Functional Coupling in Human Brain.” bioRxiv: 2024.2011. 2014.623700.

Song, D. and Z. Wang (2024). “Neurotransmitters Contribute Structure-Function Coupling: Evidence from Grey Matter Volume (GMV) and Brain Entropy (BEN).” bioRxiv: 2024.2009. 2007.611832.

Song, D. and Z. Wang (2024). “Phenotype Prediction Using BEN-Based Predictive Modeling (BPM).” bioRxiv: 2024.2009. 2007.611838.

Song, Y.-H., J. Yoon and S.-H. Lee (2021). “The role of neuropeptide somatostatin in the brain and its application in treating neurological disorders.” Experimental & Molecular Medicine 53(3): 328–338.

Sydnor, V. J., B. Larsen, D. S. Bassett, A. Alexander-Bloch, D. A. Fair, C. Liston, A. P. Mackey, M. P. Milham, A. Pines and D. R. Roalf (2021). “Neurodevelopment of the association cortices: Patterns, mechanisms, and implications for psychopathology.” Neuron 109(18): 2820–2846.

Urban-Ciecko, J. and A. L. Barth (2016). “Somatostatin-expressing neurons in cortical networks.” Nature Reviews Neuroscience 17(7): 401–409.

Van Essen, D. C., S. M. Smith, D. M. Barch, T. E. Behrens, E. Yacoub, K. Ugurbil and W.-M. H. Consortium (2013). “The WU-Minn human connectome project: an overview.” Neuroimage 80: 62–79.

Wang, Z. (2021). “The neurocognitive correlates of brain entropy estimated by resting state fMRI.” NeuroImage 232: 117893.

Wang, Z., Y. Li, A. R. Childress and J. A. Detre (2014). “Brain entropy mapping using fMRI.” PloS one 9(3): e89948.

Zhang, X.-H., K. M. Anderson, H.-M. Dong, S. Chopra, E. Dhamala, P. S. Emani, D. Margulies and A. J. Holmes (2023). “The Cellular Underpinnings of the Human Cortical Connectome.” bioRxiv.

